# Promoters adopt distinct dynamic manifestations depending on transcription factor context

**DOI:** 10.1101/650762

**Authors:** Anders S. Hansen, Christoph Zechner

## Abstract

Cells respond to external signals and stresses by activating transcription factors (TF), which induce gene expression changes. Previous work suggests that signal-specific gene expression changes are partly achieved because *different* gene promoters exhibit varying induction dynamics in response to the *same* TF input signal. Here, using high-throughput quantitative single-cell measurements and a novel statistical method, we systematically analyzed transcription in individual cells to a large number of dynamic TF inputs. In particular, we quantified the scaling behavior among different transcriptional features extracted from the measured trajectories such as the gene activation delay or duration of promoter activity. Surprisingly, we found that even the *same* gene promoter can exhibit qualitatively distinct induction and scaling behaviors when exposed to different dynamic TF contexts. That is, promoters can adopt context-dependent “manifestations”. Our analysis suggests that the full complexity of signal processing by genetic circuits may be significantly underestimated when studied in specific contexts only.

## INTRODUCTION

Exquisite regulation of gene expression underlies essentially all biological processes, including the remarkable ability of a single cell to develop into a fully formed organism. Transcription factors (TFs) control gene expression by binding to the promoters of genes and recruiting chromatin remodelers and the general transcriptional machinery. Recruitment of RNA Polymerase II enables the initiation of transcription, which produces mRNAs that are exported to the cytoplasm, where they are finally translated into proteins by the ribosome. Gene expression is primarily regulated at the level of promoter switching dynamics and initiation of transcription in a highly stochastic manner^1^. For practical reasons, however, gene expression is typically analyzed at the level of mRNAs (e.g. FISH) or proteins (e.g. immunofluorescence or GFP-reporters) using bulk or single-cell approaches. Although powerful, these data provide only partial and indirect information about the underlying promoter states and transcription initiation dynamics. Moreover, although gene regulation is complex in both time (e.g. time-varying signals) and space (e.g. signaling gradients), experimental measurements tend to be limited to simple perturbations such as ON/OFF or dose-dependent responses under steady-state conditions.

Ideally, gene regulation should be studied at the level of promoter switching dynamics and transcription initiation events, using experimental approaches that capture gene expression in a sufficiently large number of cells in response to a broad range of dynamic inputs. Several elegant studies have addressed some, but not all, of these challenges^1–5^. Here, through an integrated experimental and computational approach, we make a first attempt to realize this goal. We focus on a simple feedback-free system, where a single inducible TF activates a target gene. Surprisingly, our approach reveals that even the simplest gene networks can display complex and counter-intuitive behaviors, which cannot be explained by simple kinetic models. In particular, we show that genes exhibit “context-dependent manifestations”, such that the same gene can switch between qualitatively different kinetic behaviors depending on which dynamic input it is exposed to.

## RESULTS

### Inferring promoter induction dynamics from quantitative single-cell trajectories

To study how genes respond to complex and dynamic TF inputs, we focus on a large data-set that we previously generated (**Supplementary Fig. 1**)^4,6^. Here, addition of a small molecule causes the budding yeast TF, Msn2, to rapidly translocate to the nucleus and activate gene expression (**Fig. 1a**). Using microfluidics, rapid addition or removal of 1NM-PP1 allowed us to control both pulse length, pulse interval and pulse amplitude of the TF (fraction of Msn2 that is activated) and simultaneously measure the single-cell response of natural and mutant Msn2 target genes using fluorescent reporters^4,6–8^ (**Fig. 1a**). We note that Msn2 naturally exhibits complex signal-dependent activation dynamics^3^, that the system is not subject to feedback^3,4^, that we replaced the target gene ORF with YFP and measured the endogenous gene response^4,6^ and that the target genes are strictly Msn2-dependent^4^. Our extensive dataset contains 30 distinct dynamical Msn2 inputs for 9 genes (210 conditions) and ∼500 cells per condition, numbering more than 100,000 single-cell trajectories in total (**Fig. 1b**).

**Figure 1.**
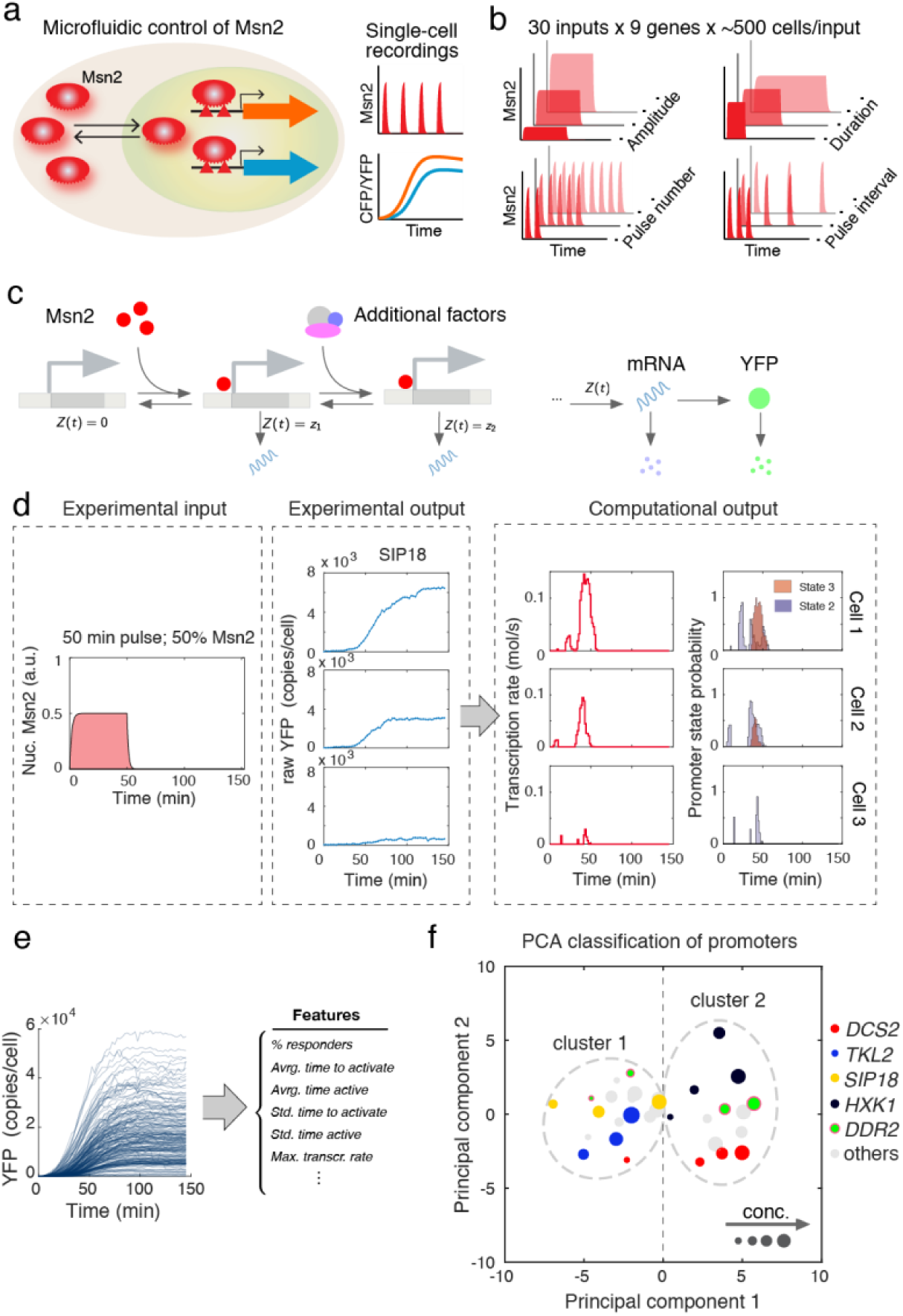
Overview of Msn2 system and inference approach. (**a**) Overview of microfluidic control of Msn2 activity and read-out of gene expression. (**b**) Overview of range of Msn2 input dynamics. **(c)** Stochastic model of gene expression. The promoter (left) can switch from its inactive state to its active state in an Msn2-dependent manner. Once active, mRNA can be transcribed at a certain rate *z*_1_. Transcription can be further tuned by recruitment of additional factors, which is captured by a third state with distinct transcription rate *z*_2_. Messenger RNA and protein dynamics are described as a two-stage birth-and-death process (right). A detailed description of the model can be found in Supplementary text S.1.1). **(d)** Statistical reconstruction of promoter switching and transcription dynamics. Gene expression output trajectories were quantified for diverse Msn2 inputs in a large number of cells. From the experimental inputs and outputs, transcription rates and promoter state occupancies were reconstructed using Bayesian inference (Box 1 and Supplementary Text S.1). **(e)** Determination of promoter features from single-cell data. Several features characterizing the promoter and transcription dynamics were calculated from the single-cell reconstructions for each promoter and experimental condition. **(f)** Clustering of promoters revealed by PCA (Supplementary text S.2.3).

Gene promoters can generally exist in a number of transcriptionally active and inactive states^9,10^. To understand the occupancy, dynamics and switching between these states, we used mathematical modeling in combination with our experimental data (**Fig. 1c**). Here we focus on a three-state promoter architecture (**Supplementary text S.1**), which was the simplest model that could recapitulate the single-cell responses of all promoters at individual Msn2 concentrations (**Supplementary text S.2.1**). This model accounts for Msn2-dependent activation of the promoter after which mRNA can be transcribed at a certain rate. Transcription can be further tuned (for instance by recruitment of additional factors), which is captured by a third state with distinct transcription rate (**Fig. 1c**).

This model in combination with a robust statistical approach allows us to determine the promoter switching- and transcription dynamics in individual cells across different promoters and conditions (see **Box 1**). However, existing inference approaches for trajectory data are limited in throughput and thus cannot handle extensive datasets like the one considered here^11–13^. We have therefore developed a novel recursive Bayesian inference method, which achieves accurate reconstructions while maintaining scalability (**Supplementary Text S.1**). The method processes single-cell trajectories and extracts from them time-varying transcription rates and promoter state occupancies (**Fig. 1d**). From the reconstructions, in turn, we computed a number of transcriptional features that summarize the single-cell expression dynamics of each promoter and condition (**Fig. 1e** and Supplementary text S.1.7). This combined experimental and computational approach allowed us to quantitatively compare different promoters under a wide range of Msn2 contexts.

To gain an overview of this high-dimensional dataset, we analyzed the gene expression responses to single pulses of nuclear Msn2 of different amplitudes (25%, 50%, 75% or 100%) for each promoter. Using Principal Component Analysis (PCA) to reduce the dimensionality (**Supplementary text S.2.3**), we uncovered the known categories of the promoters^4^ for most conditions (**Fig. 1f**): slow activation, high amplitude threshold promoters (*SIP18, TKL2*) clustered together and fast activation, low amplitude threshold promoters (*HXK1, DCS2*) also clustered together. Surprisingly, however, *DDR2* (green; **Fig. 1f**) clustered with the slow, high threshold promoters at low Msn2 amplitudes (25%, 50%), but with the fast, low threshold promoters at high Msn2 amplitudes (75%, 100%). To explain this phenomenon, we introduce the concept of “context-dependent manifestations”. Operationally, we define a context-dependent manifestation of a promoter as a situation where the same promoter exhibits qualitatively distinct kinetic behaviors under different input contexts. Here for instance, *DDR2* behaves like one promoter class at low Msn2 amplitudes, but a distinct class at high Msn2 amplitudes.

### Promoters exhibit context-dependent scaling behaviors

Having established that a single gene promoter can exhibit distinct context-dependent manifestations, we next considered specific examples of this. We analyzed the scaling behavior among different transcriptional features under all input contexts. While certain promoters behaved consistently under all contexts, others exhibited complex and context-dependent scaling behaviors. In **Fig. 2a**, for instance, we plotted the time it took to activate the promoter and the time the promoter was active (**Fig. 2b**) against the maximal transcription rate for single pulse inputs for *DDR2*. Surprisingly, at low amplitude Msn2 input, the time it takes to activate *DDR2* for the first time increases with pulse length (**Fig. 2a**) without strongly changing the active duration (**Fig. 2b**) while the maximal transcription rate remained constant (**Fig. 2a**). In contrast, at high Msn2 amplitude, the time to activate is fixed at around 5 min, but now the maximal transcription rate increases with pulse duration. Thus, we observe qualitatively distinct scaling behaviors depending on Msn2 context (here, amplitude) and refer to this phenomenon as a context-dependent manifestation.

**Figure 2:**
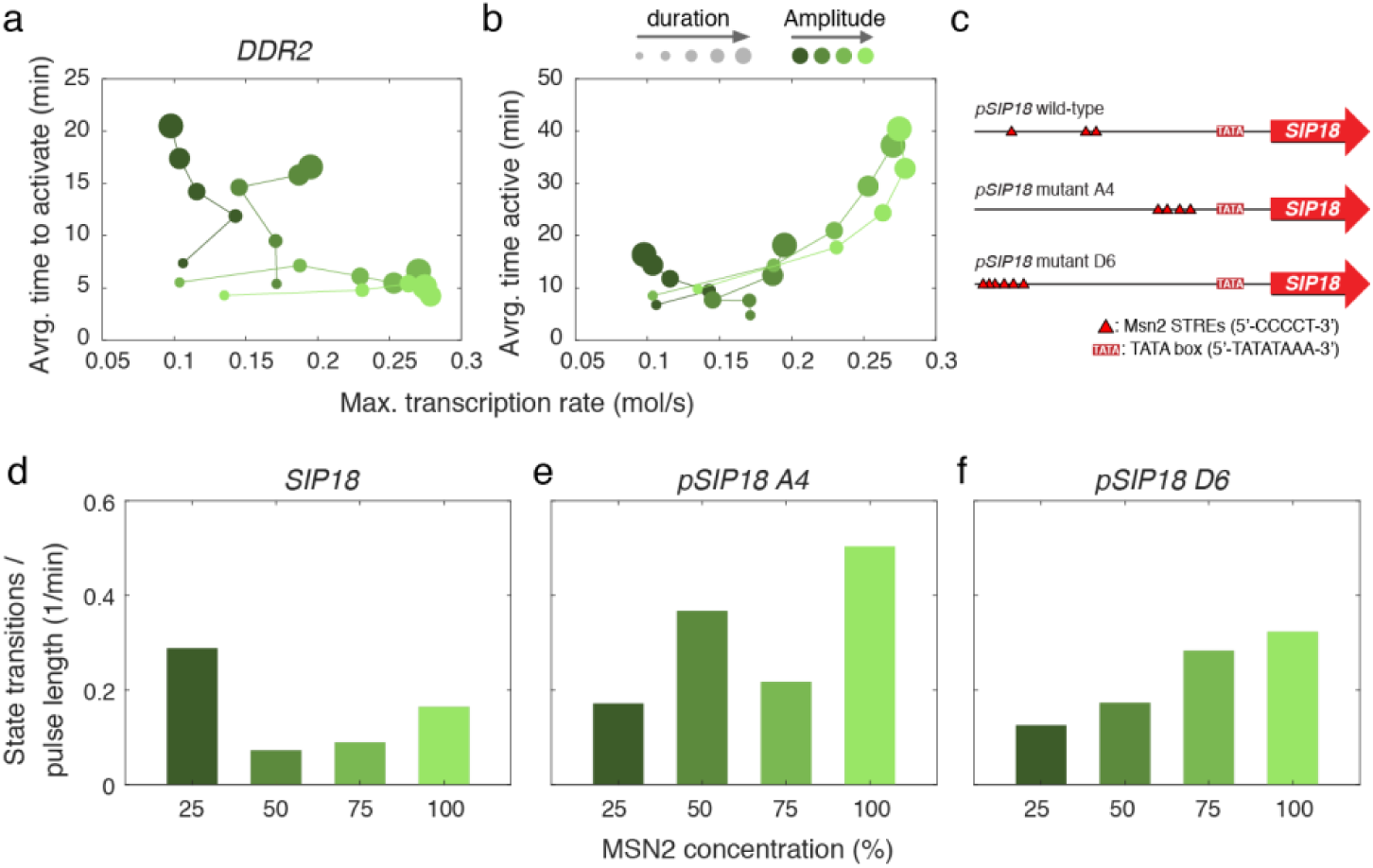
Context-dependent scaling behaviors. (**a**) Scaling behavior for *DDR2*. Scaling of time to activate (top) and total time active (bottom) for *DDR2* with the maximal transcription rate. Time to activate is defined as the time from when Msn2 enters the nucleus until the promoter converts into a transcriptionally active state. Similarly, time active is defined as the total duration the promoter is in a transcriptionally active state. Cells that never switched into a transcriptionally active state (non-responders) were excluded from the analysis. The maximal transcription rate is defined as the maximum of the average transcription rate calculated across the whole population. **(b)** Overview of wild-type and mutant *SIP18* promoters^6^. (**c-f**) Plots of the total number of state transitions normalized by pulse duration for 100% Msn2 input for wild-type (c), mutant A4 (d) and mutant D6 (f).

Having observed qualitatively distinct scaling behaviors for a natural promoter (*DDR2*), we next tested whether context-dependent behaviors are tunable by analyzing wild-type *SIP18* and two mutants^6^ of *SIP18* (**Fig. 2c**). Specifically, we analyzed the mean number of state transitions between active and inactive promoter states for a single Msn2 pulse of variable duration (10, 20, 30, 40 or 50 min) as a function of the amplitude (25%, 50%, 75% or 100% Msn2 activation) and calculated the mean number of state transitions normalized by pulse duration. Within the context of a traditional “random-telegraph” model, high transcription is normally associated with a large number of state transitions^1,4^. Surprisingly however, *SIP18* exhibited an unusual “U-shaped” scaling where low and high Msn2 amplitudes were associated with fast switching kinetics, whereas intermediate Msn2 amplitudes showed **∼**2-3 fold slower switching kinetics (**Fig. 2d**). A simple model where the switching rates increase with Msn2 amplitude cannot explain this behavior. To further investigate this scaling behavior, we compared the behavior of the wild-type *SIP18* promoter to two mutants^6^, A4 and D6 (**Fig. 2a**). While D6 (**Fig. 2f**) displayed the expected scaling behavior of monotonically increasing switching kinetics with increasing Msn2 amplitude, A4 (**Fig. 2e**) also showed unusual scaling behavior of “increasing→decreasing→increasing” switching kinetics with Msn2 concentration. In summary, these results demonstrate that a single promoter can exhibit context-dependent scaling behavior that cannot be explained by one simple model, and additionally, that only a few promoter mutations are sufficient to completely alter this behavior.

#### BOX 1. Bayesian reconstruction of promoter switching dynamics from time-lapse data.

We developed an efficient Bayesian method to quantify promoter switching and transcription dynamics from time-lapse fluorescent reporter measurements (**Fig 1, Supplementary text S.1.1**). The dynamic state of the gene expression system at time *t* is denoted by *s*(*t*) = (*z*(*t*), *m*(*t*), *n*(*t*)), with *z*(*t*) as the instantaneous transcription rate and *m*(*t*) and *n*(*t*) as the mRNA and YFP reporter copy numbers, respectively. The dynamics of *s*(*t*) are described by a continuous-time Markov chain (CTMC) as described in **Supplementary Text S.1**. Let us denote by *s*_0:*K*_ = {*s*(*t*) | 0 ≤ *t* ≤ *t*_*K*_} a complete trajectory of *s*(*t*) on the time interval *t* ∈ [0, *t*_*K*_]. We assume that we can collect a sequence of *K* partial and noisy measurements *y*_1_, …, *y*_*K*_ at times *t*_1_ < *t*_2_ < … < *t*_*K*_ along the trajectory. The statistical relationship between the measurements and the underlying state of the system is captured by a measurement density *p*(*y*_*k*_ | *s*_*k*_) with *s*_*k*_ = *s*(*t*_*k*_) for all *k* = 1, …, *K*. In the scenario considered here, the measurements *y*_1_, …, *y*_*K*_ represent noisy readouts of the reporter copy number extracted from time-lapse fluorescence movies (Materials and Methods). In order to infer *s*_0:*K*_ from a measured trajectory *y*_1_, …, *y*_*K*_, we employ Bayes’ rule, which is stated as

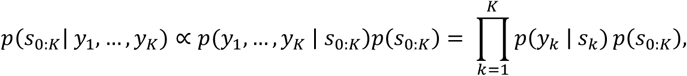

with *p*(*s*_0:*K*_) as the probability distribution over trajectories *s*_0:*K*_. The latter is governed by the CTMC model of gene expression, which summarizes the statistical knowledge about the system that is available prior to performing the experiment. The corresponding posterior distribution on the left-hand side captures the knowledge about *s*_0:*K*_ that we gain once we take into account the experimental measurements. However, analytical expressions to analyze the posterior distribution are generally lacking and one is typically left with numerical approaches. Sequential Monte Carlo (SMC) methods22 have been successfully applied to address this problem in the context of time-lapse reporter measurements^11–13^. The core idea of these approaches is to generate a sufficiently large number of random sample paths 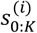 consistent with the posterior distribution from eq. (3). This is performed sequentially over individual measurement time points, which allows splitting the overall sampling problem into a sequence of smaller ones that can be solved more efficiently (**Supplementary Text S.1.4**). The resulting SMC methods, however, are still computationally very expensive since the generation of an individual sample path 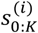 can span thousands or even millions of chemical events when considered on realistic experimental time scales. In the Msn2 induction system, for instance, trajectories involve a large number of transcription and translation events, which would render conventional SMC approaches inefficient. To address this problem, we developed a hybrid SMC algorithm, in which only the promoter switching events have to simulated explicitly, while the transcription and translation dynamics are eliminated from the simulation. More precisely, the scheme targets the marginal posterior distribution

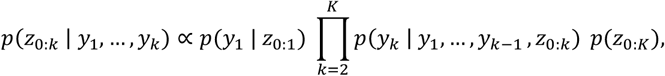

where the mRNA and reporter dynamics *m*(*t*) and *n*(*t*) have been integrated out. We derived expressions for the marginal likelihood functions *p*(*y*_*k*_ | *y*_1_, …, *y*_*k*-1_, *z*_0:*k*_) using an analytical approximation based on conditional moment equations (**Supplementary Text S.1.5**). Using this hybrid SMC algorithm, the sampling space could be significantly reduced, which makes the method scale to large datasets as the one considered in this study. Prior to applying the scheme to the dataset, the model parameters were inferred using a subset of the data (**Supplementary Text S.2.1** and **Supplementary Table 1**). A complete description of the method and additional details can be found in **Supplementary Text S.1** and **S.2**.

### Context-dependent promoter manifestations control gene expression noise

Having analyzed population-averaged properties, we next asked whether manifestations exist at the single-cell level. We began by analyzing the correlation between transcriptional output (the total number of mRNA produced) and the time the promoter was active (**Fig. 3a**) for *TKL2* and *DCS2*. For *DCS2*, transcriptional output at the single-cell level is a nearly deterministic function of the time the promoter spends in an active state. To validate this, we performed a regression analysis and found that a simple linear model where transcriptional output is proportional to time spent in the active promoter states (with coefficient β) can explain nearly all the variation in transcriptional output (*R*^2^∼1; **Fig. 3a**). Thus, for a given Msn2 amplitude, the rate of *DCS2* transcription is fixed and the single-cell expression level is determined almost exclusively by the time the promoter is active. However, the rate of transcription is set by the Msn2 amplitude (i.e. β increases with Msn2 amplitude). Thus, *DCS2* is remarkably simple and regulation by time active and transcription rate can be decoupled.

**Figure 3:**
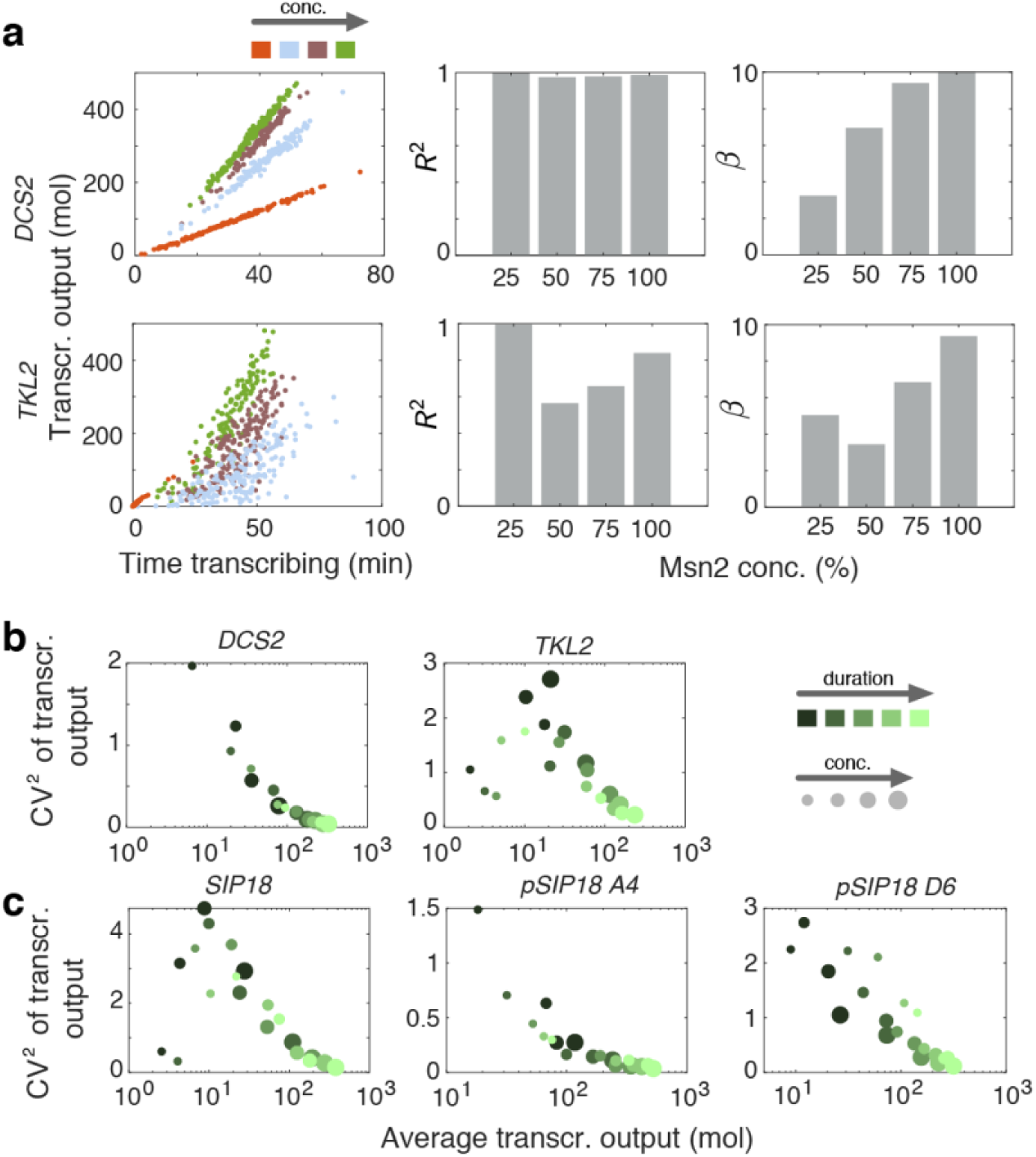
Single-cell manifestations control gene expression noise. **(a)** Scaling of transcriptional output with time transcribing, defined as the time the promoter spends in any of the two transcriptionally permissive states (see **Fig. 1c**). Transcriptional output is defined as the expected number of transcripts produced along the duration of the experiment. The left panel plots transcr. output against time transcribing for individual cells for a 50min pulse with 25%, 50%, 75% and 100% Msn2 input. Linear regression analysis was performed to determine the *R*^2^ and slope *β* between transcr. output and time transcribing as shown in the center and right panels. **(b)** Scaling of noise with average transcriptional output for different pulse lengths and Msn2 induction levels. Noise is defined as the squared coefficient variation of the transcriptional output calculated across individual cells. **(c)** Like (**b**), but for wild-type and mutant *SIP18* promoters^6^.

In contrast, *TKL2* resembles *DCS2* at low and high Msn2 amplitudes, but at 50% Msn2 activation (**Fig. 3a**, blue), *TKL2* becomes highly stochastic and time active becomes a poor predictor of transcriptional output (*R*^2^≪1; **Fig. 3a**). Thus, surprisingly, time active is an almost deterministic predictor of *TKL2* transcriptional output at low and high Msn2 amplitudes (*R*^2^∼1), but a fairly poor predictor at intermediate Msn2 amplitude (*R*^2^≪1). This demonstrates that *TKL2* exhibits an unusual “inverse-U” “deterministic→stochastic→deterministic” scaling behavior and further highlights how promoters exhibit context-dependent single-cell manifestations, in this case dependent on the Msn2 amplitude.

The analysis above considered the single-cell responses to a single 50 min Msn2 pulse at different amplitudes. To generalize our analysis, we next analyzed how transcriptional noise (quantified using CV: std/mean) scales with mean transcriptional output under all conditions (**Fig. 3b**). As expected from previous studies^14,15^, transcriptional noise uniformly decreases as transcriptional output increases for some genes such as *DCS2*. In contrast, *SIP18* exhibits the same “inverse-U” scaling as *TKL2*: low noise during low transcription, high noise during intermediate levels of transcription and again low noise during high levels of transcription (**Fig. 3c**). To further investigate the “inverse-U” scaling, we compared the behavior of the wild-type *SIP18* promoter to the two mutants A4 and D6^6^ (**Fig. 2c**). Mutant A4 behaves like *DCS2*, demonstrating that only a few promoter mutations are sufficient to switch scaling and manifestation behavior.

### Memory dependent promoter manifestations revealed by pulsatile Msn2 activation

We next analyzed how promoters respond to pulsatile Msn2 activation. Cells were exposed to four 5-min Msn2 pulses separated by 5, 7.5, 10, 15 or 20 min intervals. Some promoters behaved relatively simply, e.g. *DCS2* (**Fig. 4a**). Essentially all cells activate the *DCS2* promoter during the first pulse (**Fig. 4b**), and the promoter displays limited “positive memory” between pulses. By “positive memory”, we refer to the fact that successive pulses of Msn2 activation increase the susceptibility of the promoter to become activated and induce higher gene expression. This has also been termed the “head-start” effect^3^. In contrast, the *SIP18* mutant D6 promoter^6^ exhibited very curious behavior: at 5-min intervals (**Fig. 4b**, top row), there was significant positive memory and most cells only activated expression in the 2^nd^, 3^rd^ or 4^th^ pulse (**Fig. 4b**, top row). In contrast, with 20 min intervals, we observed “negative memory”: there was much lower expression during pulse 2-4, than during pulse 1. In other words, exposure to 1 pulse of Msn2 inhibited transcription during subsequent pulses. Furthermore, comparing the different pulse intervals we observed a transition from positive memory at 5 and 7.5 min intervals to negative memory at 15 and 20 min intervals (**Fig. 4b**). While positive memory has previously been reported^3,4^, a context-dependent switch from positive to negative memory has not. We note that no simple model can explain a sharp transition from positive to negative promoter memory and that this type of behavior only becomes visible once the response to diverse dynamic inputs are analyzed. Although the underlying molecular mechanism is unknown, we show in **Supplementary Fig. 2** a hypothetical toy model that could explain this switch from positive to negative memory. In conclusion, these data provide another example of how the same promoter can exhibit very different quantitative and qualitative behaviors depending on the context – in this case, depending on the interval between Msn2 pulses.

**Figure 4.**
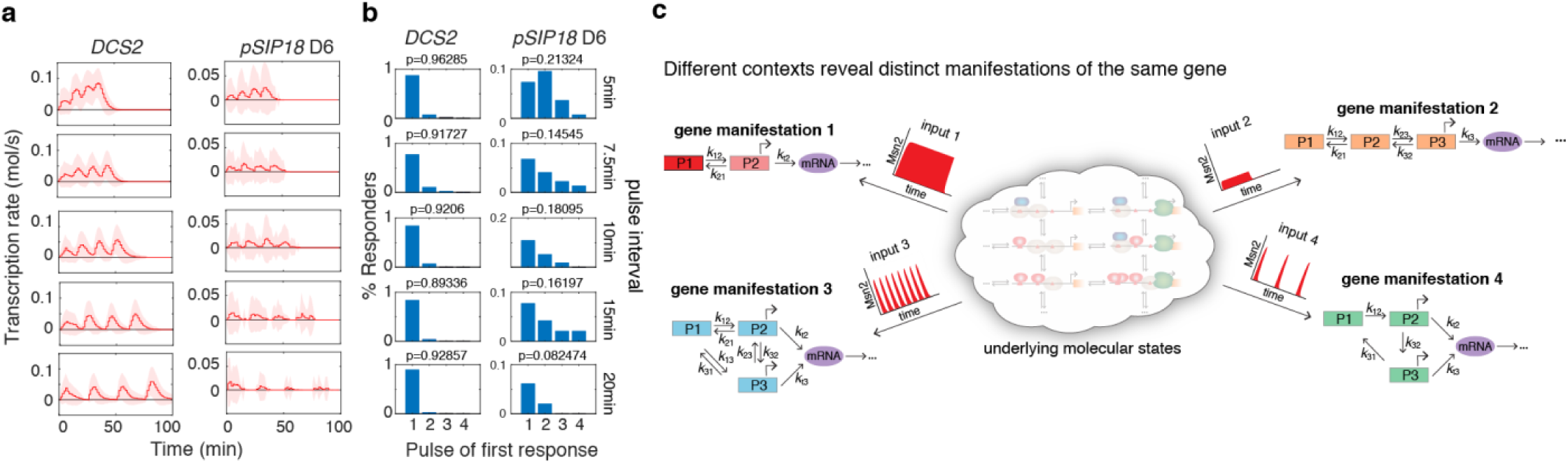
Promoter memory and model. **(a-b)** Interval-dependent regulation of promoter memory. Single-cell reconstructions are shown for *DCS2* and *SIP18* mutant D6 for the pulse-sequence experiments in which cells where treated with four consecutive Msn2 pulses (75% induction level) with 5-minute duration. The interval between the pulses were chosen to be 5, 7.5, 10, 15 and 20min. **(a)** Reconstructed transcription rates across all cells. Solid lines correspond to population means and shaded areas mark one standard deviation above and below the mean. **(b)** Distribution of responding cells across individual pulses. The histograms count for each pulse *i* the percentage of cells that had their first activation event during pulse *i*. The total percentage of responding cells *p* is provided for each histogram. **(c)** Model of context-dependent promoter manifestations.

## DISCUSSION

Here we quantitatively analyze the input-output relationship in a simple feedback-free system. We show that even under these relatively simple conditions, the same promoter can exhibit context-dependent scaling and induction behaviors. To describe this observation, we introduce the concept of context-dependent “manifestations”. The underlying number of molecular states of a promoter is potentially enormous: when we measure a dose-response, we likely observe only certain rate-limiting regimes or manifestations of the system (**Fig. 4c**). What we show here is that the particular observed rate-limiting manifestation is highly context-dependent and very distinct quantitative behaviors can be observed under different contexts – even in systems that are seemingly simple. It appears likely that many other genes and pathways exhibit similar context-dependencies, when analyzed at high resolution and under a wide range of experimental conditions.

Our results have two important implications. First, system identification efforts based on limited sets of experimental conditions within complex systems are unlikely to be successful in the sense of capturing the full range of behaviors of the underlying molecular pathways. In extreme cases, we may arrive at very different and possibly contradictory conclusions about a pathway’s inner working depending on which experimental context we choose. The only solution to this problem is to resort to experimental and computational approaches that capture a pathway’s response to a sufficiently broad range of dynamic experimental contexts. Much more work on simple systems will be necessary to truly understand the relevant complexity of signal processing in cells.

Second, a major conundrum in quantitative biology has been how to reconcile the remarkable spatiotemporal precision of biological systems with the high degree of gene expression noise observed at the single-cell level^16,17^. For example, when information transduction capacities have been measured for simple pathways, such systems appear to be barely capable of reliable distinguishing ON from OFF (∼1 bit)^18–21^. Since these studies were done under strict experimental conditions, it is likely that they captured only one out of multiple manifestations. Our results suggest that if all physiologically relevant manifestations could be captured, the estimated information transduction capacity of biochemical pathways could be substantially greater than previously estimated. This may, in part, explain the remarkable signal processing capacity of biological systems.

## MATERIALS AND METHODS; SUPPLEMENTARY FIGURES

### Overview of experiments and source data

The data in concentration units of arbitrary fluorescence were previously described^4,6^. Here, we performed absolute abundance quantification as previously described^23^ to convert the data to absolute numbers of YFP and CFP proteins per cell. All the source data supporting this manuscript are freely available together with a detailed ReadMe file at https://zenodo.org/record/2755026.

### Microfluidics and time-lapse microscope

Since the unnormalized data was previously acquired, here we only briefly describe the experimental methods. Microfluidic devices were constructed as previously described^4^. We furthermore refer the reader to a detailed protocol describing how to construct microfluidic devices and computer code for controlling the solenoid valves^8^. Briefly, for microscopy experiments, diploid yeast cells were grown overnight at 30°C with shaking at 180 RPM to an OD_600 nm_ of ca. 0.1 in low fluorescence medium without leucine and tryptophan, quickly collected by suction filtration and loaded into the five channels of a microfluidic device pretreated with concanavalin A (4 mg/mL). The setup was mounted on an inverted fluorescence microscope kept at 30°C. The microscope automatically maintains focus and acquires phase-contrast, YFP, CFP, RFP and iRFP images from each of five microfluidic channels for 64 frames with a 2.5 min time resolution corresponding to imaging from −5 min to 152.5 min. Solenoid valves control delivery of 1-NM-PP1 to each microfluidic channel. For full details on the range of input conditions, please see source data at https://zenodo.org/record/2755026.

### Image analysis and YFP quantification and normalization

Time-lapse movies were analyzed using custom-written software (MATLAB) that automatically segments yeast cells based on phase-contrast images and tracks cells between frames. The image analysis software and a protocol describing how to use it is available elsewhere^8^. The arbitrary fluorescence units were converted to absolute abundances by comparing fluorescence to strains with known absolute abundances and by segmenting the cell to calculate the total number of YFP molecules per cell per timepoint as described previously^23^.

### Quantification of nuclear Msn2 dynamics

Msn2 was visualized as an Msn2-mCherry fusion protein. This allows accurate quantification of the nuclear concentration of Msn2 over time (Msn2 only activates gene expression when nuclear) as previously described^3,4,6^. From the resulting time courses, we extracted continuous functions *u*(*t*), which served as inputs to our stochastic promoter model (**Supplementary Text S.1**). Since we found nuclear Msn2 concentration to vary very little between cells (Supplementary Figure 1), we considered *u*(*t*) to be deterministic. We performed this as described previously^4^ and elaborated on here. We model nuclear Msn2 import with first-order kinetics:

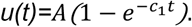

That is, if Msn2 is cytoplasmic at time *t*=0, the nuclear level of Msn2 at a later time, *t*, is given by the above expression where *A* is the maximal level of nuclear Msn2 for the given concentration of 1-NM-PP1. We chose the 1-NM-PP1 concentrations as 100 nM, 275 nM, 690 nM and 3000 nM such that they would correspond to approximately 25%,50%, 75% and 100% of maximal nuclear Msn2. *c*_1_ is a fit parameter describing the rate of nuclear import, which we found to vary slightly depending on the 1-NM-PP1 concentration.

Similarly, we model export of Msn2 from the nucleus as a first-order process:

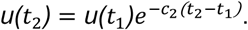

Here, *u*(*t*_*1*_) is the nuclear level of Msn2 when the microfluidic device was switched to medium without 1-NM-PP1. Correspondingly, *u*(*t*_*2*_) is the nuclear level of Msn2 at some later time *t*_2_>*t*_1._ This is to account for the fact that, depending on the pulse duration, Msn2 may not have reached its maximal nuclear level, *A*.

The parameters *A, c*_1_ and *c*_2_ were determined through fitting. Specifically, we took the full 30 different pulses and inferred the best-fit values for *A,c*_1_ and *c*_2_ using least squares fitting.

## Acknowledgements

ASH acknowledges support from Howard Hughes Medical Institute (to Erin K. O’Shea), Siebel Stem Cell Foundation (post-doctoral fellowship) and the National Institutes of Health (NIGMS K99GM130896) during parts of this work. CZ acknowledges support from the Max Planck Society, the MPI-CBG as well as the Center for Advancing Electronics Dresden (cfaed). We thank Nan Hao, Nadine Vastenhouw, Stephan Grill, Andre Nadler, Carl Modes, Alf Honigmann, Pavel Tomancak and Tony Hyman for insightful comments on the manuscript.

---

**Supplementary Figure 1.**
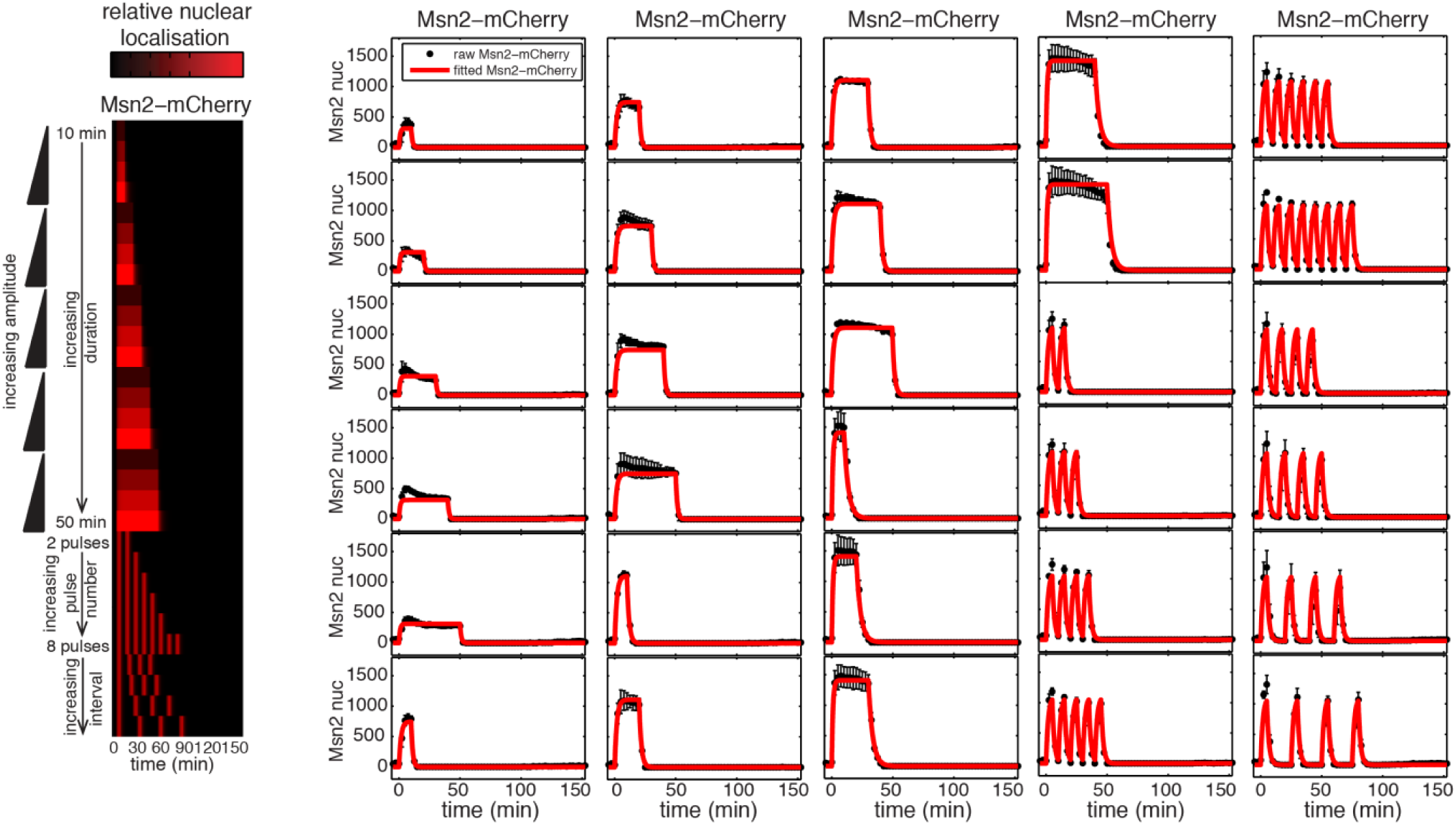
Overview of Msn2 input experiments. Left: heatmap overview of the 30 different Msn2-mCherry input. Right: Raw experimentally measured Msn2-mCherry input (black) and standard deviation (black error bars) for each of the 30 Msn2 inputs. The fitted Msn2-mCherry input is overlaid in red. This figure has been partially reproduced with permission from *Molecular Systems Biology*^4^.

**Supplementary Figure 2.**
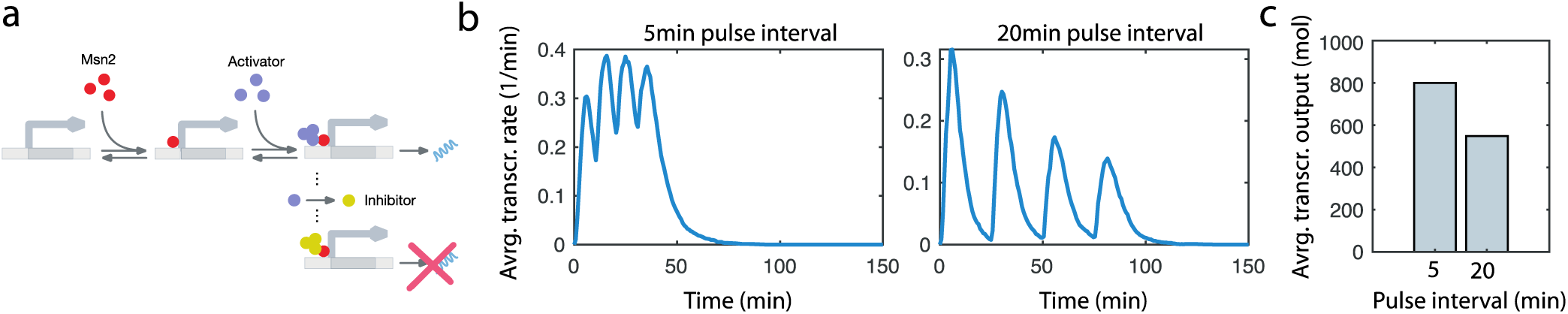
Toy model with interval-dependent promoter memory. (**a**) Model scheme. Once Msn2 binds to the promoter, activator molecules can be recruited, which causes the promoter to switch into a transcriptionally active state with a rate proportional to the number of activators present. Once the promoter switches back into the Msn2-unbound state, the activator can be converted into an inhibitor, which causes the promoter to switch into a transcriptionally inactive state with a rate proportional to the number of inhibitors present. (**b**) Average transcription rate for 5 min and 20 min pulse intervals as a function of time obtained by forward simulation of the model. Blue lines indicate averages computed from stochastic simulations (n=2000). (**c**) Corresponding average transcriptional output for 5min and 20min pulse intervals. A detailed reaction scheme and parameters used for simulation can be found in Supplementary text S.3. We emphasize that this Toy model only serves to illustrate one possible scenario, which could result in a pulse-interval dependent switch from positive to negative memory, as observed in **Fig. 4a-b**. We do not currently understand the mechanism underlying the observation in **Fig. 4a-b**.

