## Supplementary material for "Promoters adopt distinct dynamic manifestations depending on transcription factor context": SI

### S.1 Statistical inference of transcription dynamics from time-course reporter measurements

#### S.1.1 Stochastic model of Msn2-dependent gene expression

We describe Msn2-dependent gene expression using a canonical three-state model as shown in Figure S.1. The promoter is described as a continuous-time Markov chain, which switches stochastically between three states of different transcriptional activity (Figure S.1a). Correspondingly, the rate of transcription at time $t$ is governed by a stochastic process $Z(t) \in \{z_0, z_1, z_2\}$, whose value changes discontinuously whenever the promoter transitions from one state into another. In the absence of nuclear Msn2, the promoter is in its transcriptionally inactive state ($z_0 = 0$), where no transcripts are produced. Upon recruitment of Msn2 to the promoter, it can switch into a transcriptionally permissive state in which transcription takes place with propensity $z_1$. To account for Msn2-dependent promoter activation, we consider the switching rate from $z_0$ to $z_1$ to depend on the nuclear Msn2 concentration. For simplicity, we consider a linear dependency, i.e., $q_{01}(t) = \gamma u(t)$, with $u(t)$ as the Msn2 abundance at time $t$. The corresponding reverse rate $q_{10}$ is considered to be constant. We assume that transcription can be further enhanced by recruitment of additional factors such as chromatin remodeling complexes and general transcriptional factors. This is captured in our model by introducing a third state with transcription rate $z_2$ and corresponding transition rates $q_{12}$ and $q_{21}$. With this, we can describe the time-dependent probability distribution over the transcription rate $P_Z(t) = (P(Z(t) = z_0 \mid \theta), P(Z(t) = z_1 \mid \theta), P(Z(t) = z_2 \mid \theta))^T$ in terms of a forward equation

$$\frac{\mathrm{d}}{\mathrm{d}t} P_Z(t) = QP(t) = \begin{pmatrix} -q_{01}(t) & q_{10} & 0 \\ q_{01}(t) & -q_{10} - q_{12} & q_{21} \\ 0 & q_{12} & -q_{21} \end{pmatrix} P_Z(t), \tag{1}$$

with $P_Z(0) = p_{z,0}$ as some initial distribution over $Z(t)$ and $\theta = \{\gamma, q_{10}, q_{12}, q_{21}\}$ as a set of parameters. In the following, we denote by $\mathbf{z}_t = \{z(s) \mid 0 \leq s \leq T\}$ a complete realization of $Z(t)$ on a fixed time interval $[0, t]$. Furthermore, we introduce the conditional path distribution $p(\mathbf{z}_t \mid \theta)$ which measures the likelihood of observing a particular trajectory $\mathbf{z}_t$ for a given parameter set $\theta$. Note that it is straightforward to draw random sample paths $\mathbf{z}_t$ from this distribution using Gillespie's stochastic simulation algorithm [2] or its variants.

Transcription and translation are modeled as a two-stage reaction network as shown in Figure S.1b. We denote by $M(t)$ and $N(t)$ the copy numbers of mRNA and protein at time $t$, respectively. The parameters $c_1$ and $c_2$ are the mRNA and protein degradation rates and $A$ is the protein translation rate. To account for cell-to-cell variability in protein translation, we consider the latter to be randomly distributed across isogenic cells, i.e., $A \sim p(a \mid \beta)$, with $p(a \mid \beta)$ as an arbitrary probability density function (pdf) with positive support and $\beta$ as a set of hyperparameters characterizing this distribution [3, 4]. Here we consider as hyperparameters the average and coefficient of variation (CV) of $A$ such that $\beta = \{\langle A \rangle, CV[A]\}$. Consequently, $\beta$ captures the magnitude and variability associated with protein translation. In the following, we denote by $\omega = \{c_1, c_2, \beta\}$ the set of parameters corresponding to transcription and translation.

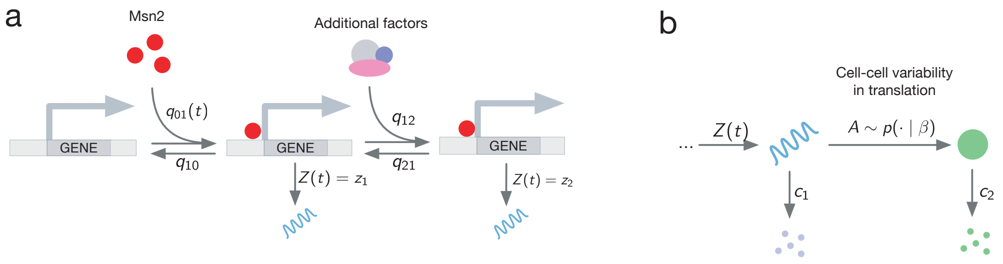


Figure S.1: Stochastic model of Msn2-inducible reporter expression. (a) Promoter model. (b) Model of transcription and translation.

For a given set of parameters $\theta$ and $\omega$ and a concrete realization of the translation rate $A$, the overall dynamics of the joint system state $(Z(t), M(t), N(t))$ can be described by a Markov chain. However, due to the random variability over $A$, each cell is associated with a differently parameterized Markov chain. This results in a so-called mixed-effect Markov model, whose computational analysis turns out to be challenging [3]. One way to address this issue is to augment the state space by the random variable $A$ and to formulate a master equation on this extended space. For $S(t) = (Z(t), M(t), N(t), A)$, such master equation reads

$$\begin{aligned}
\frac{\mathrm{d}}{\mathrm{d}t} P(z_0, m, n, a, t) = {} & z_0 P(z_0, m-1, n, a, t) + c_1(m+1)P(z_0, m+1, n, a, t) \\
& + amP(z_0, m, n-1, a, t) + c_2(n+1)P(z_0, m, n+1, a, t) \\
& - [z_0 + c_1 m + am + c_2 n]\, P(z_0, m, n, a, t) \\
& + q_{10} P(z_1, m, n, a, t) - q_{01}(t) P(z_0, m, n, a, t) \\
\frac{\mathrm{d}}{\mathrm{d}t} P(z_1, m, n, a, t) = {} & z_1 P(z_1, m-1, n, a, t) + c_1(m+1)P(z_1, m+1, n, a, t) \\
& + amP(z_1, m, n-1, a, t) + c_2(n+1)P(z_1, m, n+1, a, t) \\
& - [z_1 + c_1 m + am + c_2 n]\, P(z_1, m, n, a, t) \\
& + q_{01}(t) P(z_0, m, n, a, t) - q_{10}(t) P(z_1, m, n, a, t) \\
& + q_{21} P(z_2, m, n, a, t) - q_{12}(t) P(z_1, m, n, a, t) \\
\frac{\mathrm{d}}{\mathrm{d}t} P(z_2, m, n, a, t) = {} & z_2 P(z_2, m-1, n, a, t) + c_1(m+1)P(z_2, m+1, n, a, t) \\
& + amP(z_2, m, n-1, a, t) + c_2(n+1)P(z_2, m, n+1, a, t) \\
& - [z_2 + c_1 m + am + c_2 n]\, P(z_2, m, n, a, t) \\
& + q_{12}(t) P(z_1, m, n, a, t) - q_{21}(t) P(z_2, m, n, a, t)
\end{aligned} \tag{2}$$

with $P(z_i, m, n, a) := P(Z(t) = z_i, M(t) = m, N(t) = n, A \in [a + \mathrm{d}a) \mid \theta, \omega)$. Differential equations for arbitrary moments $\mathbb{E}\left[f(Z(t), M(t), N(t), A)\right]$ with $f$ as a polynomial can be computed by multiplying (3) with $f$ and summing or integrating over all possible values of $m$, $n$, $z_i$ and $a$, respectively [4]. Since the network is linear in its propensities, the resulting moment equations turn out to be closed and can thus be computed without approximation. Moreover, we will denote by $\mathbf{s}_t = \{s(u) \mid 0 \leq u \leq t\}$ a complete sample path of the full system state between time zero and $t$.

### S.1.2 Conditional dynamics of transcription and translation

One major difficulty in inferring gene networks like the one in Figure S.1 is that they involve both very lowly and highly abundant components. This is why moment-based descriptions of the full system state $S(t)$ are of limited use for the time-series inference problem considered here as will be discussed later in Section S.1.3. On the other hand, approaches purely based on stochastic simulation become computationally expensive, since transcription and translation often involve thousands or even millions of events over the duration of a time-course experiment. In such cases, hybrid approaches can be beneficial, where only the lowly abundant components are described stochastically, whereas the remaining components are handled using moment equations. In the scenario considered here, for instance, the time evolution of the transcription rate $\mathbf{z}_t$ can be efficiently simulated using stochastic simulation since the number of times the promoter switches between states is comparably small. For a given $\mathbf{z}_t$, one could then calculate a corresponding set of conditional moments characterizing the dynamics of mRNA and protein. More technically, this can be understood by the fact that the path distribution over the total system state factorizes into $p(\mathbf{s}_t \mid \omega, \theta) = p(\mathbf{x}_t \mid \mathbf{z}_t, \omega)p(\mathbf{z}_t \mid \theta)$. Correspondingly, we can describe the dynamics over $X(t) = (M(t), N(t), A)$ as a conditional Markov process $X(t) \mid \mathbf{z}_t$, whose state probability distribution $P(m, n, a, t) := P(M(t) = m, N(t) = n, A \in [a + \mathrm{d}a) \mid \mathbf{z}_t)$ satisfies

$$\begin{aligned}\frac{\mathrm{d}}{\mathrm{d}t}P(m,n,a,t) = {} & z(t)P(m-1,n,a,t) + c_1(m+1)P(m+1,n,a,t) \\ & + amP(m,n-1,a,t) + c_2(n+1)P(m,n+1,a,t) \\ & - [z(t) + c_1 m + am + c_2 n]\, P(m,n,a,t),\end{aligned} \tag{3}$$

whereas we assume for the initial condition $P(m, n, a, t = 0) = P(M(0) = m, N(0) = n \mid Z(0) = z_0)p(a \mid \beta)$. In order the derive conditional moments $\mathbb{E}[X(t) \mid \mathbf{z}_t]$, we multiply (3) with polynomials in $x$ and sum and integrate over all $m$, $n$ and $a$, respectively. Here, we consider moments of mRNA and protein up to order two, which can be fully described by the system of differential equations

$$\begin{aligned}
&\frac{\mathrm{d}}{\mathrm{d}t}\mathbb{E}[M(t) \mid \mathbf{z}_t] = z(t) - \mathbb{E}[M(t) \mid \mathbf{z}_t]\, c_1 \\
&\frac{\mathrm{d}}{\mathrm{d}t}\mathbb{E}[N(t) \mid \mathbf{z}_t] = \mathbb{E}[M(t)A \mid \mathbf{z}_t] - \mathbb{E}[N(t) \mid \mathbf{z}_t]\, c_2 \\
&\frac{\mathrm{d}}{\mathrm{d}t}\mathbb{E}\left[M(t)^2 \mid \mathbf{z}_t\right] = z(t) + 2\mathbb{E}[M(t) \mid \mathbf{z}_t]\, z(t) + \mathbb{E}[M(t) \mid \mathbf{z}_t]\, c_1 - 2\mathbb{E}\left[M(t)^2 \mid \mathbf{z}_t\right] c_1 \\
&\frac{\mathrm{d}}{\mathrm{d}t}\mathbb{E}[M(t)N(t) \mid \mathbf{z}_t] = \mathbb{E}[N(t) \mid \mathbf{z}_t]\, z(t) - \mathbb{E}[M(t)N(t) \mid \mathbf{z}_t]\, c_1 - \mathbb{E}[M(t)N(t) \mid \mathbf{z}_t]\, c_2 + \mathbb{E}\left[M(t)^2 A \mid \mathbf{z}_t\right] \\
&\frac{\mathrm{d}}{\mathrm{d}t}\mathbb{E}[M(t)A \mid \mathbf{z}_t] = \mathbb{E}[A \mid \mathbf{z}_t]\, z(t) - \mathbb{E}[M(t)A \mid \mathbf{z}_t]\, c_1 \\
&\frac{\mathrm{d}}{\mathrm{d}t}\mathbb{E}\left[N(t)^2 \mid \mathbf{z}_t\right] = \mathbb{E}[N(t) \mid \mathbf{z}_t]\, c_2 + \mathbb{E}[M(t)A \mid \mathbf{z}_t] - 2\mathbb{E}\left[N(t)^2 \mid \mathbf{z}_t\right] c_2 + 2\mathbb{E}[M(t)N(t)A \mid \mathbf{z}_t] \\
&\frac{\mathrm{d}}{\mathrm{d}t}\mathbb{E}[N(t)A \mid \mathbf{z}_t] = \mathbb{E}\left[M(t)A^2 \mid \mathbf{z}_t\right] - \mathbb{E}[N(t)A \mid \mathbf{z}_t]\, c_2 \\
&\frac{\mathrm{d}}{\mathrm{d}t}\mathbb{E}\left[M(t)^2 A \mid \mathbf{z}_t\right] = \mathbb{E}[A \mid \mathbf{z}_t]\, z(t) + 2\mathbb{E}[M(t)A \mid \mathbf{z}_t]\, z(t) + \mathbb{E}[M(t)A \mid \mathbf{z}_t]\, c_1 - 2\mathbb{E}\left[M(t)^2 A \mid \mathbf{z}_t\right] c_1 \\
&\frac{\mathrm{d}}{\mathrm{d}t}\mathbb{E}[M(t)N(t)A \mid \mathbf{z}_t] = \mathbb{E}[N(t)A \mid \mathbf{z}_t]\, z(t) - \mathbb{E}[M(t)N(t)A \mid \mathbf{z}_t]\, c_1 - \mathbb{E}[M(t)N(t)A \mid \mathbf{z}_t]\, c_2 \\
&\qquad + \mathbb{E}\left[M(t)^2 A^2 \mid \mathbf{z}_t\right] \\
&\frac{\mathrm{d}}{\mathrm{d}t}\mathbb{E}\left[M(t)A^2 \mid \mathbf{z}_t\right] = \mathbb{E}\left[A^2 \mid \mathbf{z}_t\right] z(t) - \mathbb{E}\left[M(t)A^2 \mid \mathbf{z}_t\right] c_1 \\
&\frac{\mathrm{d}}{\mathrm{d}t}\mathbb{E}\left[M(t)^2 A^2 \mid \mathbf{z}_t\right] = \mathbb{E}\left[A^2 \mid \mathbf{z}_t\right] z(t) + 2\mathbb{E}\left[M(t)A^2 \mid \mathbf{z}_t\right] z(t) + \mathbb{E}\left[M(t)A^2 \mid \mathbf{z}_t\right] c_1 \\
&\qquad - 2\mathbb{E}\left[M(t)^2 A^2 \mid \mathbf{z}_t\right] c_1.
\end{aligned} \tag{4}$$

Note that (4) involves all first and second order moments, but also a few additional moments of order three and four, which are needed in order to obtain a closed set of differential equations.

#### S.1.3 Statistical model of time-series reporter measurements

As detailed in Materials and Methods, we performed quantitative single-cell time-lapse measurements of reporter expression for different Msn2-inducible promoters and Msn2 activation profiles. We denote by $t_1, \ldots, t_K$ the time points at which measurements were taken. Correspondingly, we define by $\mathbf{s}_{l:k}$ a complete sample path of the gene expression system between times $t_l$ and $t_k$. If $l = 0$, we refer the state at time $t = 0$, which does not necessarily coincide with the first measurement time point $t_1$. The measurements – denoted by $Y_k$ for $k = 1, \ldots, K$ – provide noisy information about the system state $S(t_k)$ according to a measurement density

$$Y_k \mid (S(t_k) = s_k) \sim p(\cdot \mid s_k).$$

We consider the measurement noise to be independent among time-points such that

$$p(y_1, \ldots, y_K \mid s_1, \ldots, s_K) = \prod_{k=1}^{K} p(y_k \mid s_k). \tag{5}$$

In our particular case, the measurements correspond to the reporter abundance $N(t)$ corrupted by measurement noise such that

$$p(y_k \mid s_k) = p(y_k \mid x_k) = p(y_k \mid n_k).$$

For a given set of parameters $\{\theta, \omega\}$, the relation between a complete sample path $\mathbf{s}_{0:K}$ and the observed measurements is captured by a joint distribution

$$p\left(y_1, \ldots, y_K, \mathbf{s}_{0:K} \mid \omega, \theta\right) = P(s_0) p(\mathbf{s}_{0:K} \mid \omega, \theta) \prod_{k=1}^{K} p(y_k \mid s_k), \tag{6}$$

with $P(s_0) := P(S(0) = s_0)$ as the initial distribution over the system state and $p(\mathbf{s}_{0:K} \mid \omega, \theta)$ as the distribution over complete sample paths $\mathbf{s}_{0:K}$. Correspondingly, the posterior distribution over $\mathbf{s}_{0:K}$ is proportional to (6), i.e.,

$$p\left(\mathbf{s}_{0:K} \mid y_1, \ldots, y_K, \omega, \theta\right) \propto P(s_0) p(\mathbf{s}_{0:K} \mid \omega, \theta) \prod_{k=1}^{K} p(y_k \mid s_k). \tag{7}$$

#### S.1.4 Recursive Bayesian Estimation

The posterior distribution (7) is generally intractable but several approximate techniques have been proposed previously [3]. Most of them rely on Bayesian filtering methods, which construct an approximation of (7) recursively over measurement time points. In those approaches, one exploits the fact that the posterior distribution at any measurement time $t_k$ can be written recursively as

$$p\left(\mathbf{s}_{0:k} \mid y_1, \ldots, y_k, \omega, \theta\right) \propto p(y_k \mid s_k) p(\mathbf{s}_{k-1:k} \mid s_{k-1}, \omega, \theta) p\left(\mathbf{s}_{0:k-1} \mid y_1, \ldots, y_{k-1}, \omega, \theta\right), \tag{8}$$

with $p\left(\mathbf{s}_{0:k-1} \mid y_1, \ldots, y_{k-1}, \omega, \theta\right)$ as the posterior distribution at time $t_{k-1}$. In order to solve the Bayesian recursion between consecutive time steps, one can either employ analytical approximations, or Monte Carlo methods. In a recent study, for instance, we have proposed a Gaussian approximation of the Bayesian filtering problem, which relies on the time-evolution of the first and second order moments of the gene network dynamics [5]. While computationally efficient, the underlying Gaussian approximation is not suitable for lowly abundant, switch-like components, such as the transcription rate $Z(t)$ in our promoter model. Alternative approaches are mostly based on sequential Monte Carlo techniques [6, 3], which approximate (8) using a sufficiently large number of Monte Carlo samples drawn by SSA. The main advantage of these techniques is that they are exact up to sampling variance but on their downside, suffer from limited scalability. In particular, forward-simulation via SSA can become prohibitively slow, especially when RNAs and proteins are highly abundant. They are therefore not able to tackle large datasets like the one considered here. In the following, we will present a novel hybrid inference algorithm, which bypasses expensive SSA simulations of highly abundant species, making it sufficiently scalable to deal with datasets that span tens- or even hundreds of thousands of single-cell trajectories.

#### S.1.5 Hybrid sequential Monte Carlo

One strategy to improve the scalability of sequential Monte Carlo techniques is to analytically eliminate variables that are not of direct interest to a particular inference problem [7, 3]. In our case, for instance, we are specifically interested in the promoter switching dynamics and the corresponding transcription rate $Z(t)$. From this perspective, it would therefore suffice to calculate the marginal posterior distribution

$$p(\mathbf{z}_{0:K} \mid y_1, \ldots, y_K, \theta, \omega) \propto p(y_1, \ldots, y_K, \mathbf{z}_{0:K} \mid \theta, \omega) = \mathbb{E}\left[p(y_1, \ldots, y_K, \mathbf{x}_{0:K}, \mathbf{z}_{0:K} \mid \theta, \omega)\right], \tag{9}$$

in which the dynamics of $X(t)$ have been "integrated out". In order to perform this integration, we first realize that the joint distribution can be rewritten as

$$\begin{aligned} p\left(y_1, \ldots, y_K, \mathbf{s}_{0:K} \mid \omega, \theta\right) &= p\left(y_1, \ldots, y_K, \mathbf{x}_{0:K}, \mathbf{z}_{0:K} \mid \omega, \theta\right) \\ &= P(s_0) p(\mathbf{s}_{0:K} \mid \omega, \theta) \prod_{k=1}^{K} p(y_k \mid s_k) \\ &= P(x_0, z_0) p(\mathbf{x}_{0:K} \mid \mathbf{z}_{0:K}, \omega) p(\mathbf{z}_{0:K} \mid \theta) \prod_{k=1}^{K} p(y_k \mid x_k) \\ &= P(x_0, z_0) p(\mathbf{z}_{0:K} \mid \theta) \prod_{k=1}^{K} p(y_k \mid x_k) p(\mathbf{x}_{k-1:k} \mid x_{k-1}, \mathbf{z}_{k-1:k}, \omega), \end{aligned} \tag{10}$$

where we have made use of the fact that $p(\mathbf{s}_{0:K} \mid \omega, \theta) = p(\mathbf{x}_{0:K} \mid \mathbf{z}_{0:K}, \omega) p(\mathbf{z}_{0:K} \mid \theta)$ as shown in Section S.1.2. Next, we integrate (10) over all subpaths $\{\mathbf{x}_{k-1:k} \setminus x_k\}$ such that only the values of $X(t)$ at the time points $t_0, \ldots, t_K$ remain in the model. This integration is straightforward and can be carried out by replacing the path distribution $p(\mathbf{x}_{k-1:k} \mid x_{k-1}, \mathbf{z}_{k-1:k}, \omega)$ by the state transition kernel $P(x_k \mid x_{k-1}, \mathbf{z}_{k-1:k}, \omega)$, i.e.,

$$p\left(y_1, \ldots, y_K, x_0, \ldots, x_K, \mathbf{z}_{0:K} \mid \omega, \theta\right) = P(x_0, z_0) p(\mathbf{z}_{0:K} \mid \theta) \prod_{k=1}^{K} p(y_k \mid x_k) P(x_k \mid x_{k-1}, \mathbf{z}_{k-1:k}, \omega). \tag{11}$$

The marginalization over the remaining variables $x_0, \ldots, x_K$ then reduces to a summation

$$p(y_1, \ldots, y_K, \mathbf{z}_{0:K} \mid \theta, \omega) = \sum_{x_0} \cdots \sum_{x_K} p\left(y_1, \ldots, y_K, x_0, \ldots, x_K, \mathbf{z}_{0:K} \mid \omega, \theta\right). \tag{12}$$

Most conveniently, this summation can be solved iteratively, by first summing over $x_0$, subsequently over $x_1$ and so forth. The first summation yields

$$\begin{aligned} p\left(y_1, \ldots, y_K, x_1, \ldots, x_K, \mathbf{z}_{0:K} \mid \omega, \theta\right) &= \sum_{x_0} P(x_0 \mid z_0) p(y_1 \mid x_1) P(x_1 \mid x_0, \mathbf{z}_{0:1}, \omega) p(\mathbf{z}_{0:K} \mid \theta) \\ &\quad \times \prod_{k=2}^{K} p(y_k \mid x_k) P(x_k \mid x_{k-1}, \mathbf{z}_{k-1:k}, \omega) \\ &= p(y_1 \mid x_1) P(x_1 \mid \mathbf{z}_{0:1}, \omega) p(\mathbf{z}_{0:K} \mid \theta) \times \prod_{k=2}^{K} p(y_k \mid x_k) P(x_k \mid x_{k-1}, \mathbf{z}_{k-1:k}, \omega) \\ &= p(\mathbf{z}_{0:K} \mid \theta) P(x_1 \mid y_1, \mathbf{z}_{0:1}, \omega) p(y_1 \mid \mathbf{z}_{0:1}, \omega) \times \prod_{k=2}^{K} p(y_k \mid x_k) P(x_k \mid x_{k-1}, \mathbf{z}_{k-1:k}, \omega), \end{aligned} \tag{13}$$

whereas the last step follows from the fact that $p(y_1 \mid x_1)P(x_1 \mid \mathbf{z}_{0:1}, \omega) = P(x_1 \mid y_1, \mathbf{z}_{0:1}, \omega)p(y_1 \mid \mathbf{z}_{0:1}, \omega)$ via Bayes' rule. Repeating the same procedure for $x_1$ yields

$$
\begin{aligned}
&p\left(y_1, \ldots, y_K, x_2, \ldots, x_K, \mathbf{z}_{0:K} \mid \omega, \theta\right) \\
&\quad = \sum_{x_1} p(\mathbf{z}_{0:K} \mid \theta) P(x_1 \mid y_1, \mathbf{z}_{0:1}, \omega) p(y_1 \mid \mathbf{z}_{0:1}, \omega) \times \prod_{k=2}^{K} p(y_k \mid x_k) P(x_k \mid x_{k-1}, \mathbf{z}_{k-1:k}, \omega) \\
&\quad = p(\mathbf{z}_{0:K} \mid \theta) p(y_2 \mid x_2) \sum_{x_1} P(x_2 \mid x_1, \mathbf{z}_{1:2}, \omega) P(x_1 \mid y_1, \mathbf{z}_{0:1}, \omega) p(y_1 \mid \mathbf{z}_{0:1}, \omega) \\
&\qquad \times \prod_{k=3}^{K} p(y_k \mid x_k) P(x_k \mid x_{k-1}, \mathbf{z}_{k-1:k}, \omega) \\
&\quad = p(\mathbf{z}_{0:K} \mid \theta) p(y_2 \mid x_2) P(x_2 \mid y_1, \mathbf{z}_{0:2}, \omega) p(y_1 \mid \mathbf{z}_{0:1}, \omega) \times \prod_{k=3}^{K} p(y_k \mid x_k) P(x_k \mid x_{k-1}, \mathbf{z}_{k-1:k}, \omega) \\
&\quad = p(\mathbf{z}_{0:K} \mid \theta) P(x_2 \mid y_2, y_1, \mathbf{z}_{0:2}, \omega) p(y_2 \mid y_1, \mathbf{z}_{0:2}, \omega) p(y_1 \mid \mathbf{z}_{0:1}, \omega) \times \prod_{k=3}^{K} p(y_k \mid x_k) P(x_k \mid x_{k-1}, \mathbf{z}_{k-1:k}, \omega).
\end{aligned}
\tag{14}
$$

Continuing the above procedure for $x_2, \ldots, x_K$ finally leads to

$$
p\left(y_1, \ldots, y_K, \mathbf{z}_{0:K} \mid \omega, \theta\right) = p(\mathbf{z}_{0:K} \mid \theta) p(y_1 \mid \mathbf{z}_{0:1}, \omega) \prod_{k=2}^{K} p(y_k \mid y_{k-1}, \ldots, y_1, \mathbf{z}_{0:k}, \omega). \tag{15}
$$

Therefore, the marginal posterior distribution over the transcription dynamics $\mathbf{z}_{0:K}$ is proportional to (15), which can again be expressed recursively as

$$
\begin{aligned}
&p\left(\mathbf{z}_{0:K} \mid y_1, \ldots, y_K, \omega, \theta\right) \propto p\left(y_1, \ldots, y_K, \mathbf{z}_{0:K} \mid \omega, \theta\right) \\
&\qquad \propto p(y_K \mid y_{K-1}, \ldots, y_1, \mathbf{z}_{0:K}, \omega) p(\mathbf{z}_{K-1:K} \mid z_{K-1}, \theta) p\left(\mathbf{z}_{0:K-1} \mid y_1, \ldots, y_{K-1}, \omega, \theta\right).
\end{aligned}
\tag{16}
$$

Importantly, using eq. (16) we can perform a sequential Monte Carlo algorithm on a significantly reduced sampling space, where only the transcription dynamics $\mathbf{z}_{0:K}$ have to be simulated explicitly. However, in order to perform this algorithm, we need to be able to calculate the marginal likelihood terms $p(y_k \mid y_{k-1}, \ldots, y_1, \mathbf{z}_{0:k}, \omega)$, which are given by

$$
\begin{aligned}
p(y_k \mid y_{k-1}, \ldots, y_1, \mathbf{z}_{0:k}, \omega) &= \sum_{x_k} p(y_k \mid x_k) P(x_k \mid y_{k-1}, \ldots, y_1, \mathbf{z}_{0:k}, \omega) \\
&= \sum_{x_k} p(y_k \mid x_k) \sum_{x_{k-1}} P(x_k \mid x_{k-1}, \mathbf{z}_{k-1:k}, \omega) P(x_{k-1} \mid y_{k-1}, \ldots, y_1, \mathbf{z}_{0:k-1}, \omega).
\end{aligned}
\tag{17}
$$

The two sums in (17) are generally intractable, but analytical solutions exist if the measurement likelihood function $p(y_k \mid x_k)$ and the state transition kernel $P(x_k \mid x_{k-1}, \mathbf{z}_{k-1:k}, \omega)$ belong to certain classes of distributions. This is the case, for instance, if both are Gaussian. However, this is likely not a good assumption in the scenario considered here, since both the measurement- and state distributions are generally positive and asymmetric. As it turns out, however, eq. (17) has an analytical solution also if both $p(y_k \mid x_k)$ and $P(x_k \mid x_{k-1}, \mathbf{z}_{k-1:k}, \omega)$ are log-normally distributed, which is in good agreement with previous studies[8, 3, 9] and also the experimental data considered here. We therefore assume

$$
\begin{aligned}
Y_k \mid (N(t) = n) &\sim \mathcal{LN}\left(\log(n), \eta^2\right) \\
X(t) \mid \mathbf{z}_t &\sim \mathcal{LN}\left(\mu(t), \Sigma(t)\right),
\end{aligned}
\tag{18}
$$

where $\eta^2$ corresponds to the strength of the measurement noise and $\mu(t) \in \mathbb{R}^3$ and $\Sigma(t) \in \mathbb{R}^{3\times 3}$ characterize the distribution over $X(t) = (M(t), N(t), A)$ conditionally on a particular realization of $\mathbf{z}_{0:K}$. More precisely, $\mu(t)$ and $\Sigma(t)$ are the mean and covariance of $\log X(t)$ and we therefore refer to them as logarithmic moments in the following.

Now, assuming that the posterior distribution over $X(t)$ is log-normally distributed at time $t_{k-1}$,

$$p(x_{k-1} \mid y_{k-1}, \dots, y_1, \mathbf{z}_{0:k-1}, \omega) \approx \mathcal{LN}\left(x_{k-1} \mid \mu(t_{k-1}), \Sigma(t_{k-1})\right), \tag{19}$$

it will – based on our assumption – remain log-normal upon applying the state transition kernel, i.e.,

$$\begin{aligned} p(x_k \mid y_{k-1}, \dots, y_1, \mathbf{z}_{0:k}, \omega) &\approx \int p(x_k \mid x_{k-1}, \mathbf{z}_{k-1:k}, \omega) \mathcal{LN}\left(x_{k-1} \mid \mu(t_{k-1}), \Sigma(t_{k-1})\right) \mathrm{d}x_{k-1} \\ &= \mathcal{LN}\left(x_k \mid \mu(t_k), \Sigma(t_k)\right), \end{aligned} \tag{20}$$

whereas the sum has now been replaced by an integral. In order to calculate the logarithmic moments $\mu(t_k)$ and $\Sigma(t_k)$ for a given $\mu(t_{k-1})$ and $\Sigma(t_{k-1})$, one first has to calculate the first and second order moments from the log-normal distribution, propagate those forward in time until $t_k$ using the conditional moment dynamics described in Section S.1.2 and subsequently convert them back into the logarithmic domain to obtain $\mu(t_k)$ and $\Sigma(t_k)$. The relationship between logarithmic and standard moments is given by

$$\begin{aligned} \mathbb{E}\left[X_i(t)\right] &= e^{\mu_i(t) + \frac{1}{2}\Sigma_{ii}(t)} \\ \mathbb{E}\left[X_i(t) X_j(t)\right] &= e^{\mu_i(t) + \mu_j(t) + \frac{1}{2}\left(\Sigma_{ii}(t) + 2\Sigma_{ij}(t) + 2\Sigma_{ji}(t) + \Sigma_{jj}(t)\right)}. \end{aligned} \tag{21}$$

In order to determine the posterior distribution at the next measurement time $t_k$, we multiply (20) with the log-normal measurement density such that

$$\begin{aligned} p(x_k \mid y_k, \dots, y_1, \mathbf{z}_{0:k}, \omega) &\propto p(y_k \mid x_k) p(x_k \mid y_{k-1}, \dots, y_1, \mathbf{z}_{0:k}, \omega) \\ &= \mathcal{LN}\left(y_k \mid \log(n_k), \eta^2\right) \times \mathcal{LN}\left(x_k \mid \mu(t_k), \Sigma(t_k)\right), \end{aligned} \tag{22}$$

with $n_k = N(t_k)$ as the protein abundance at time $t_k$. It is straightforward to show that the product of two log-normal distributions in (22) is again a log-normal distribution such that

$$p(x_k \mid y_k, \dots, y_1, \mathbf{z}_{0:k}, \omega) = \mathcal{LN}\left(x_k \mid \mu^+(t_k), \Sigma^+(t_k)\right), \tag{23}$$

with

$$\Sigma^+(t_k) = \left[\frac{1}{\eta^2} w w^T + \Sigma(t_k)^{-1}\right]^{-1} \tag{24}$$

$$\mu^+(t_k) = \Sigma^+(t_k) \left[\frac{1}{\eta^2} \log(y_k) w + \Sigma(t_k)^{-1} \mu(t_k)\right], \tag{25}$$

with $w = (0, 1, 0)^T$ as a vector that reflects the fact that from $X(t) = (M(t), N(t), A)$, the second component (i.e., the protein abundance) is measured experimentally.

For the likelihood term $p(y_k \mid y_{k-1}, \dots, y_1, \mathbf{z}_{0:k}, \omega)$ we obtain

$$\begin{aligned} p(y_k \mid y_{k-1}, \dots, y_1, \mathbf{z}_{0:k}, \omega) &= \int p(y_k \mid x_k) p(x_k \mid y_{k-1}, \dots, y_1, \mathbf{z}_{0:k}, \omega) \mathrm{d}x_k \\ &= \int \mathcal{LN}\left(y_k \mid \log(n_k), \eta^2\right) p(n_k \mid y_{k-1}, \dots, y_1, \mathbf{z}_{0:k}, \omega) \mathrm{d}n_k \\ &= \int \mathcal{LN}\left(y_k \mid \log(n_k), \eta^2\right) \mathcal{LN}\left(n_k \mid \mu_2(t_k), \Sigma_{22}(t_k)\right) \mathrm{d}n_k, \end{aligned} \tag{26}$$

whereas the last line follows from the fact that each dimension $i$ of a multivariate log-normal distribution with logarithmic moments $\mu$ and $\Sigma$ is marginally log-normal with parameters $\mu_i$ and $\Sigma_{ii}$. This integral can be solved in closed form such that we obtain for the logarithm of the marginal likelihood function

$$\begin{aligned}\log p(y_k \mid y_{k-1}, \dots, y_1, \mathbf{z}_{0:k}, \omega) = \\ -\frac{1}{2}\left[\frac{(\log y_k - \mu_2(t_k))^2}{\eta^2 + \Sigma_{22}(t_k)} - \log\left(\frac{1}{\eta^2} + \Sigma_{22}(t_k)^{-1}\right) - \log \Sigma_{22}(t_k) - \log \eta^2\right] + const.\end{aligned} \tag{27}$$

Together, eqs. (4), (24), (25) and (27) define a recursive Bayesian filter, which allows us to eliminate the components $X(t)$ from the inference problem. As mentioned above, the remaining component $Z(t)$ can then be inferred efficiently using a conventional sequential importance sampler. To this end, we define a set of $J$ particles, each of them consisting of a path $\mathbf{z}^{(i)}$, a set of logarithmic moments $\mu^{(i)}$ and $\Sigma^{(i)}$ as well as a particle probability $p^{(i)}$. This set of particles serves as a finite sample approximation of the posterior distribution at each iteration $k$. At the $k$th time step, $J$ new particles are drawn randomly according to the particle probabilities $p^{(i)}$. For each particle $i$, the path $\mathbf{z}^{(i)}$ is first extended to the next measurement $t_{k+1}$ using SSA. The new probability of this particle is then determined by first propagating the corresponding logarithmic moments until $t_{k+1}$ using eq. (4) and then evaluating eq. (27). The particle probabilities are then normalized across the $J$ particles such that they sum up to one. Subsequently, $\mu^{(i)}$ and $\Sigma^{(i)}$ are updated using (24) and (25) and the algorithm proceeds with the next iteration. At the final time $t_K$, the paths $\mathbf{z}^{(i)}$ associated with the particles represent samples from the desired marginal posterior distribution, which can be used for further analysis.

#### S.1.6 Quantitative characterization of promoter dynamics

The inference algorithm described above allows as to compute an arbitrary number of samples $\mathbf{z}_{0:K}^{(i)}$ from the desired posterior distribution. In order to compare the dynamics of the different promoters under various experimental conditions, we extracted a number of features from these samples that characterize the transcriptional response for each individual cell. More technically, these features can be defined as functionals that map a random path $\mathbf{z}_{0:K}^{(i)}$ to a real or discrete number. This functional can then be averaged with respect to the posterior distribution associated with a particular cell, i.e.,

$$\mathbb{E}\left[f(\mathbf{z}_{0:K}) \mid y_1, \dots, y_K\right] \approx \frac{1}{J}\sum_{j=1}^{J} f(\mathbf{z}_{0:K}^{(j)}), \tag{28}$$

with $y_1, \dots, y_K$ as the measurements of this cell and $\mathbf{z}_{0:K}^{(j)}$ as samples from the posterior distribution obtained from the inference method. The following list summarizes the different features that were used in this study.

- **Percentage of responders.** A cell is considered a responder if it managed to switch into a state of significant transcriptional activity at least once. To this end, we defined a functional $r(\mathbf{z}_{0:K}) \in \{0, 1\}$, which is one only if a promoter state was reached which had a transcription rate of at least 20% of the maximum transcription rate taken over all Msn2 induction levels. Depending on the promoter and condition, this could encompass one, two or none of the promoter states. We then estimated the response probability $p_a = \mathbb{E}\left[r(\mathbf{z}_{0:K}) \mid y_1, \dots, y_K\right]$ for each cell by averaging over all the individual samples paths $\mathbf{z}_{0:K}$ obtained from the sequential Monte Carlo algorithm. A cell was then classified as a responder if $p_a > 0.99$. Subsequently, we calculated the percentage of responders for each promoter and condition.
- **Time to activate.** For all responding cells, we calculated the posterior expectation of the time it took until the cell switched into a transcriptionally significant state, i.e., $\mathbb{E}\left[\tau_S(\mathbf{z}_{0:K}) \mid y_1, \dots, y_K\right]$ with $\tau_S(\mathbf{z}_{0:K}) \in \mathbb{R}^+$ as a functional that measures the time until the first transition into a responsive state happened. We further calculated the mean and variance of the time until activation over all cells in an experiment.

- **Total time active.** Analogously to the time to activate, we quantified the total time the promoter was active, i.e., $\mathbb{E}\left[\tau_A(\mathbf{z}_{0:K}) \mid y_1, \ldots, y_K\right]$ with $\tau_S(\mathbf{z}_{0:K}) \in \mathbb{R}^+$ as a functional that extracts the total time the promoter spent in any of the active states.
- **Time spent in state $i$.** We calculated the total time the promoter spent in any of the three states, i.e., $\mathbb{E}\left[\tau_i(\mathbf{z}_{0:K}) \mid y_1, \ldots, y_K\right]$ with $\tau_i(\mathbf{z}_{0:K}) \in \mathbb{R}^+$.
- **Number of transitions from state $i$ to $j$.** For each cell we calculated how often the promoter switched between states 1 and 2, and 2 and 3, respectively. More technically, we estimated the expectation $\mathbb{E}\left[n_{ij}(\mathbf{z}_{0:K}) \mid y_1, \ldots, y_K\right]$ with $n_{ij}(\mathbf{z}_{0:K}) \in \mathbb{N}$ as a functional that counts the number of transitions from state $i$ to $j$. Additionally, we counted the total number of transitions that took place over the duration of the time course experiment.
- **Maximum transcription.** We calculated the maximum transcription rate that the promoter achieved during a time course experiment. In particular, we computed the expected transcription rate for each cell $\lambda(t) = \mathbb{E}\left[Z(t) \mid y_1, \ldots, y_K\right]$ and subsequently the corresponding population average $\langle\lambda(t)\rangle$. We then determined the maximum of this average, i.e, $\lambda_{max} = \max_t \langle\lambda(t)\rangle$.
- **Time to maximum transcription.** Next to the maximum transcription, we also determined the time when this maximum was achieved, i.e., $\tau_{max} = \arg\max_t \langle\lambda(t)\rangle$.

#### S.1.7 Evaluation of the inference method using synthetic data

In order to study the accuracy of the proposed inference method, we tested it using artificially generated data. In particular, we considered two differently parameterized versions of the stochastic model in Figure S.1. The first one resembled a fast promoter like *DCS2* or *HXK1* whereas the second one had slow and switch-like promoter activation kinetics like *SIP18* or *TKL1*. In particular, the parameters of the system were chosen to be $\gamma = 0.05$, $q_{10} = 0.055$, $q_{12} = 0.001\kappa$, $q_{21} = 0.004\kappa$, $z_1 = 0.0035$, $z_2 = 0.728$, $c_1 = 0.0013$, $c_2 = 1.67e-5$, $\langle A\rangle = 0.1$, $CV[A] = 0.02$, whereas $\kappa = \{1, 10\}$ for the slow and fast promoter model, respectively. All rate parameters are given in units $s^{-1}$.

For each promoter, we generated 30 single cell trajectories between time zero and $t_K = 150\,\text{min}$ using SSA and sampled the protein abundance at 55 equidistant time points $t_1, \ldots, t_K$. For the Msn2 activation function $u(t)$, we used the experimentally determined profile for a single pulse experiment (75% Msn2 induction level, 40min duration). The measurements were then simulated from a log-normal measurement density $\mathcal{LN}\left(y_k \mid \log(n_k), \eta^2\right)$, with $n_k$ as the protein copy number at time $t_k$ and $\eta$ as the logarithmic standard deviation of this density. For this study, we set $\eta = 0.05$.

We applied the hybrid sequential Monte Carlo algorithm to reconstruct the promoter dynamics and compared it to the true realization. In particular, we analyzed three of the path functionals described in Section S.1.6: total time active, time to activate and transcriptional output. We estimated posterior expectations of these functionals using $J = 400$ Monte Carlo samples and analyzed how they compared to the true values extracted from the exact sample paths $\mathbf{z}_{0:K}$. We first assumed perfect knowledge of all process parameters. The top panels in Figure S.2a and b show the inferred values plotted against the ground truth. For all three features we found a strict linear relationship with a slope close to one. Moreover, the corresponding high $R^2$ values show that the inference method operates at a very high accuracy. We furthermore analyzed the robustness of the method with respect to parameter mismatch. To this end, we randomly perturbed all of the parameters using a log-normal distribution $\mathcal{LN}\left(\log a, 0.1^2\right)$ with $a$ as the underlying true value. Note that this was performed for each of the considered trajectories separately. In case of poor robustness, we would thus expect a significantly reduced correlation. However, we found for all three features that both the $R^2$ and slope changed only marginally indicating a high robustness of the method. This is an important feature in practical scenarios where knowledge about process parameters is generally imperfect.

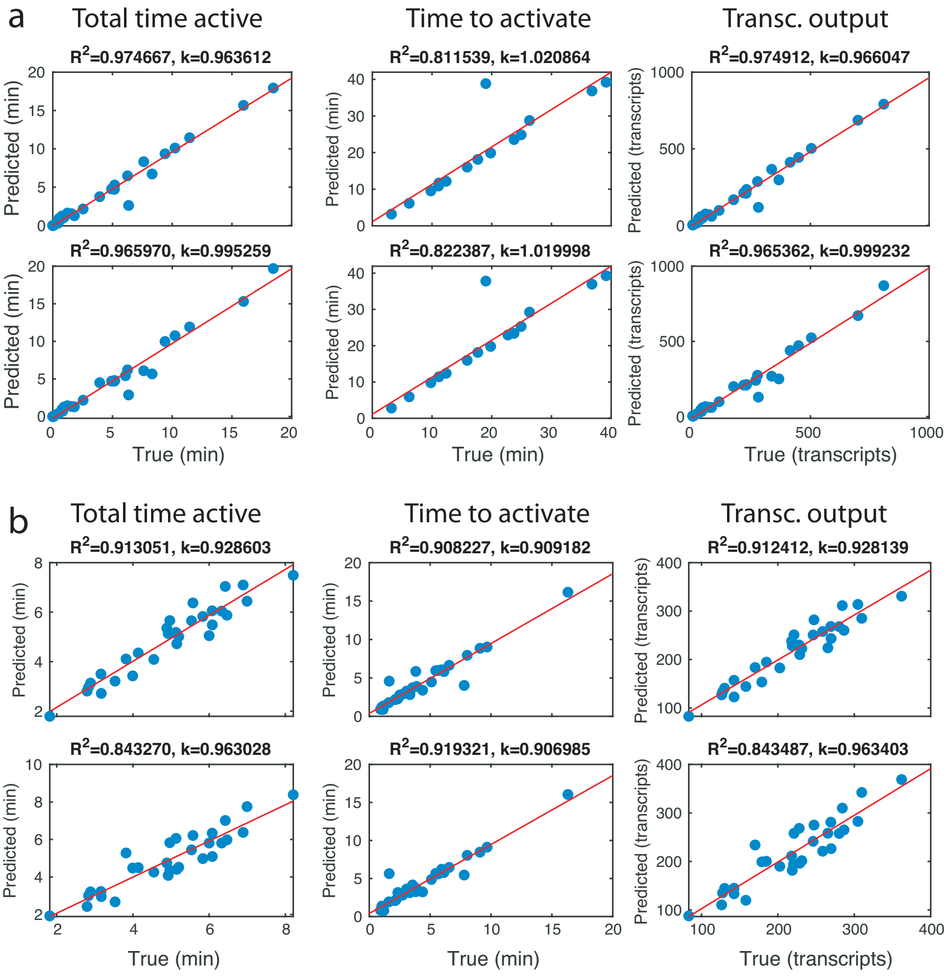


Figure S.2: Evaluation of the inference method using synthetic data. The inference method was performed using artificially generated time-course data as described in the text. (a) Inference results for the slow promoter model. (b) Inference results for the fast promoter model. Respective top panels show the results assuming perfect knowledge of the model parameters. Bottom panels show the corresponding results under parameter mismatch. The $R^2$ and slope $k$ between the true and predicted features were determined using linear regression (red lines).

### S.2 Statistical analysis of Msn2-dependent gene expression

In the following we provide details on the statistical analysis of Msn2-dependent gene expression as shown in the main text. In this case, the function $u(t)$ corresponds to the nuclear Msn2 concentration that was measured experimentally for each condition (Materials and Methods and Supplementary Figure 1). In combination with the measured YFP time series, this allowed us to infer and quantitate the input-output relationship of different promoters under different experimental conditions using the recursive inference method described in Section S.1.4. However, before this method could be applied, the stochastic model from Figure S.1 had to be parameterized. To for this purpose, we used a portion of the experimental single-cell trajectories to infer the kinetic parameters of the model (Section S.2.1). Subsequently, we reconstructed the transcription dynamics of each promoter and condition as described in Section S.2.2.

#### S.2.1 Statistical inference of kinetic parameters

In order to parameterize the stochastic gene expression model for different promoters and experimental conditions, we used an established moment-based inference method [4]. This method uses a Markov chain Monte Carlo sampler to match the first and second order moments of the stochastic gene network to the experimentally determined ones. For detailed information on this approach, the reader shall refer to [4]. The kinetics of the same gene expression system have been previously quantified using a simpler, deterministic model [10]. We incorporated this additional information in the form of suitable prior distributions over some of the kinetic parameters. In particular, we considered Gamma prior distributions $p(c_1) = \Gamma(20, 20/1.3e-3s^{-1})$ and $p(\langle A \rangle) = \Gamma(20, 20/0.05s^{-1})$ for the mRNA degradation and average protein translation rates, respectively. Additionally, the protein degradation rate was fixed to $c_2 = 1.67e-5s^{-1}$. For the switching parameters $q_{ij}$, we used a prior distribution $p(q_{ij}) = \Gamma(1, 1/30s^{-1})$. For each promoter, we first estimated the total set of parameters $\omega$ and $\theta$ using the single pulse experiments with maximum level and duration (100% Msn2, 50min). In particular, we used a Metropolis-Hastings sampler with lognormal proposal distributions to generate $2e4$ samples, whereas the first $3e3$ samples were discarded. From the resulting samples we extracted maximum a posterior (MAP) estimates of the model parameters. Since the promoter switching dynamics is generally concentration-dependent, we re-estimated $\theta$ for the 25%, 50% and 75% Msn2 pulse experiments (also 50min pulse duration). However, we expected the transcriptional and translational dynamics to remain the same among different experiments of the same promoter and we thus kept $\omega$ fixed across different Msn2 induction levels. Maximum a posterior (MAP) estimates of the parameters for the different promoters and conditions are summarized in Supplementary Table 1. The resulting model fits are shown Figure S.3.

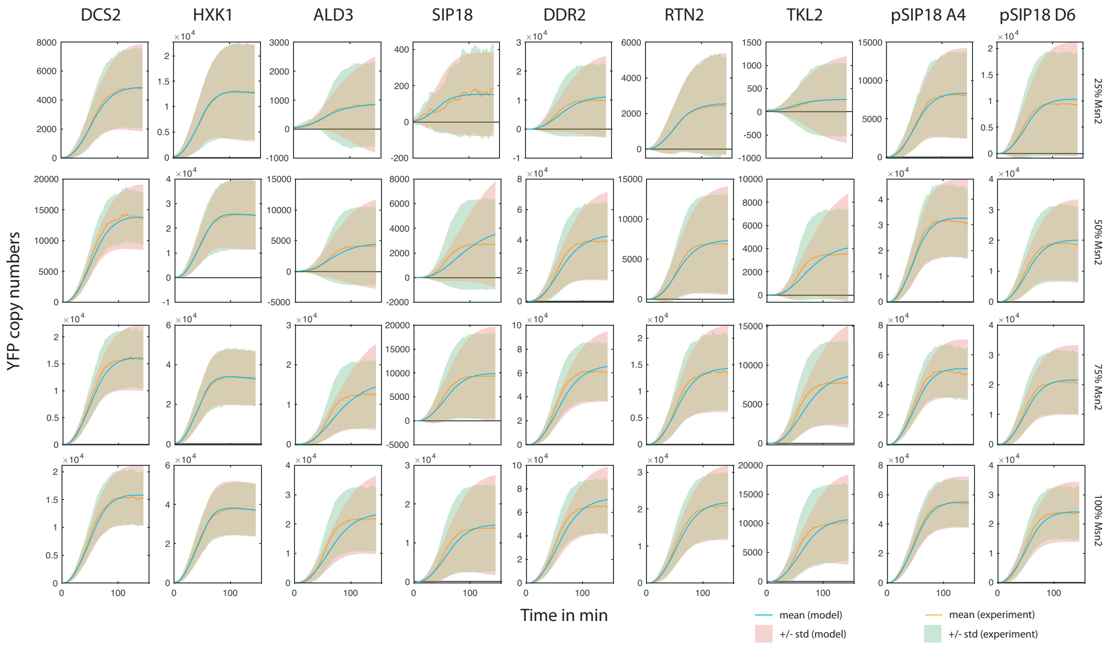


Figure S.3: Comparison of the calibrated model with the experimental data.

#### S.2.2 Statistical inference of transcription dynamics

Based on the calibrated models, we inferred the transcription and promoter switching dynamics using the recursive Bayesian inference scheme from Section S.1.4. Based on our previous study [3] which uses a similar data processing and calibration pipeline, we set the measurement noise parameter to $\eta = 0.15$ corresponding to an expected relative variation of around 15 percent. For each condition and promoter, we processed each individual cell using $J = 400$ particles. From the resulting samples, we calculated Monte Carlo estimates of the promoter features summarized in Section S.1.6.

#### S.2.3 Clustering of promoters

To get an overview of how promoters behave under different contexts, we performed a principal component analysis (PCA) of the previously inferred promoter features. For this purpose, we considered all single pulse experiments (10-50min duration, 25-100% Msn2 induction). For each condition, we calculated the percentage of responders, the average time to activate, the average time active and the maximal transcription rate. For each promoter and Msn2 concentration, we concatenated the respective features for all pulse lengths, giving rise to a 20-dimensional feature vector. Each individual feature (e.g., average time active) was normalized across all five pulse lengths and promoters to preserve the relative scaling of this feature with pulse length. In total this leads to 36 20-dimensional data points (4 concentrations for 9 promoters), which we analyzed using PCA. Figure 1f in the main text shows the first two leading principal components plotted against each other.

### S.3 Toy model of interval-dependent promoter memory

For the simulations shown in Supplementary Fig. 2, we considered a simple promoter model described by a reaction network

$$\begin{aligned} P_0 &\underset{c_2}{\overset{c_1 u(t)}{\rightleftharpoons}} P_1 \\ P_1 &\underset{c_4}{\overset{c_3 A(t)}{\rightleftharpoons}} P_2 \\ P_0 &\underset{c_6}{\overset{c_5 I(t)}{\rightleftharpoons}} P_3 \\ P_1 &\xrightarrow{c_7} P_1 + A \\ P_0 + A &\xrightarrow{c_8} P_0 + I, \end{aligned} \tag{29}$$

with $u(t)$ as the experimentally measured nuclear Msn2 abundance (Materials and Methods). Transcription takes place with rate $z$ when the promoter is in state $P_2$. The parameters used for simulation were chosen to be $c_1 = 0.02$, $c_2 = 0.06$, $c_3 = 0.003$, $c_4 = 0.02$, $c_5 = 0.0006$, $c_6 = 0.001$, $c_7 = 0.9$, $c_8 = 7e - 6$ and $z = 0.01$ in units $s^{-1}$.
